# Co-evolutionary analysis reveals a conserved dual binding interface between extracytoplasmic function (ECF) σ factors and class I anti-σ factors

**DOI:** 10.1101/2020.04.09.035246

**Authors:** Delia Casas-Pastor, Angelika Diehl, Georg Fritz

## Abstract

Extracytoplasmic function σ factors (ECFs) belong to the most abundant signal transduction mechanisms in bacteria. Amongst the diverse regulators of ECF activity, class I anti-σ factors are the most important signal transducers in response to internal and external stress conditions. Despite the conserved secondary structure of the class I anti-σ factor domain (ASDI) that binds and inhibits the ECF under non-inducing conditions, the binding interface between ECFs and ASDIs is surprisingly variable between the published co-crystal structures. In this work, we provide a comprehensive computational analysis of the ASDI protein family and study the different contact themes between ECFs and ASDIs. To this end, we harness the co-evolution of these diverse protein families and predict covarying amino acid residues as likely candidates of an interaction interface. As a result, we find two common binding interfaces linking the first α-helix of the ASDI to the DNA binding region in the σ_4_ domain of the ECF, and the fourth α-helix of the ASDI to the RNA polymerase (RNAP) binding region of the σ_2_ domain. The conservation of these two binding interfaces contrasts with the apparent quaternary structure diversity of the ECF/ASDI complexes, partially explaining the high specificity between cognate ECF and ASDI pairs. Furthermore, we suggest that the dual inhibition of RNAP- and DNA-binding interfaces are likely a universal feature of other ECF anti-σ factors, preventing the formation of non-functional trimeric complexes between σ/anti-σ factors and RNAP or DNA.

**Significance:** In the bacterial world, extracytoplasmic function σ factors (ECFs) are the most widespread family of alternative σ factors, mediating many cellular responses to environmental cues, such as stress. This work uses a computational approach to investigate how these σ factors interact with class I anti-σ factors – the most abundant regulators of ECF activity. By comprehensively classifying the anti-σs into phylogenetic groups and by comparing this phylogeny to the one of the cognate ECFs, the study shows how these protein families have co-evolved to maintain their interaction over evolutionary time. These results shed light on the common contact residues that link ECFs and anti-σs in different phylogenetic families and set the basis for the rational design of anti-σs to specifically target certain ECFs. This will help to prevent the cross-talk between heterologous ECF/anti-σ pairs, allowing their use as orthogonal regulators for the construction of genetic circuits in synthetic biology.

## Introduction

Extracytoplasmic function σ factors (ECFs) are one the most abundant signal transduction mechanisms in the bacterial kingdom, often mediating the cellular response to external and internal stress conditions. Although these minimalistic members of the σ^70^ family contain only the σ_2_ and σ_4_ domains essential for recruiting RNA polymerase (RNAP) to specific promoter sequences (1), ECFs have evolved in into a surprisingly diverse protein family. By now we know 156 phylogenetic ECF groups (2, 3) – many of which feature group-specific target promoter motifs and conserved regulators of ECF activity, suggesting similar modes of signal transduction within an ECF group. Amongst these diverse signaling mechanisms, the most common regulators of ECF activity are so-called anti-σ factors, which, under non-inducing conditions, sequester ECF into inactive complexes via their anti-σ domain (ASD). Under inducing conditions, anti-σ factors releases their ECFs by various mechanisms, including anti-σ proteolysis (4–6), conformational change (7, 8) or sequestration by ECF-mimicking anti-anti-σ factors (9, 10). Given that bacteria harbor an average of 10 ECFs, and some species encode more than 100 ECFs per genome (3), the pertinent question arises how the different ECF/anti-σ factor pairs prevent massive cross-talk between each other, or in other words, how they achieve signaling specificity?

Here we focus on the class I anti-σ factors, which are not only the first characterized, but also the most abundant anti-σs known to date (2, 11). Class I anti-σ factors are defined by their N-terminal anti-σ domain I (ASDI), which features a common secondary structure consisting of four alpha helices: The first three (N-terminal) helices form a bundle that binds to the σ_4_ domain of the ECF and, separated by a flexible linker, the fourth helix binds to the σ_2_ domain. Interestingly, while this general theme has been found in all of the four crystal structures of ASDI/ECF complexes solved to date (6, 12–14), these structures also expose a significant diversity in the binding topology between ECFs and ASDIs (Fig. 1). The most striking difference relates to the overall ECF/ASDI conformation (Fig. 1), showing that in three of the co-crystal structures, ChrR/SigE_*Rsp*_ (*Rhodobacter sphaeroides*), RseA/RpoE_*Eco*_ (*Escherichia coli*) and RskA/SigK_*Mtu*_ (*Mycobacterium tuberculosis*), the ASDI is sandwiched between the σ_2_ and σ_4_ domains, while RsiW wraps around the two σ domains of SigW (*Bacillus subtilis*). Furthermore, while the three-helix bundle of some ASDI require zinc coordination for ECF inhibition (ChrR_*Rsp*_; (11)), another structure features a zinc-binding motif but binds the ECF independent of zinc (RsiW_*B. subtilis*_; (12)), and others do not rely on a zinc binding motif at all (RskA_*Mtu*_, RseA_*Eco*_; (13, 14)). Thus, it is tempting to speculate that the divergent binding topologies between ECFs and ASDIs could be important to prevent cross-talk between ECF/ASDI pairs of different ECF groups, and that these conformations might be conserved within ECF groups. If so, we reasoned that protein sequences of ECF and ASDI proteins have co-evolved, and that ASDI protein sequences should cluster into phylogenetic groups similar to the ECF groups.

**Figure 1.**
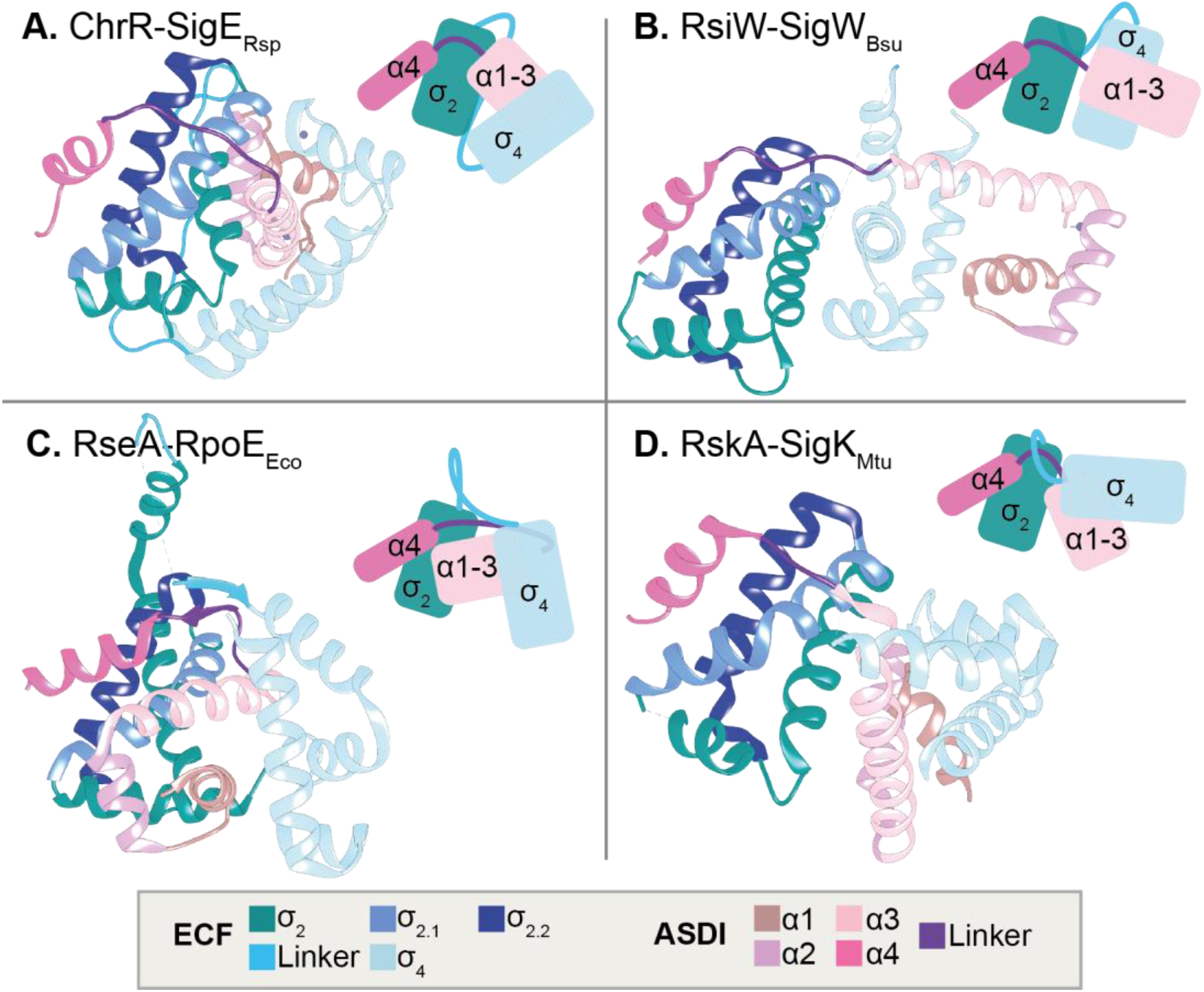
Structures of ECF σ factors in complex with class I anti-σ factors. ECFs are shown in pink colors, whereas anti-σ factors appear in blue colors. Different areas of the protein are differentially colored (see legend). Different anti-σ factors show different binding conformations. **A**: SigE-ChrR from *R. sphaeroides* (PDB: 2Q1Z (11)). **B**: SigW-RsiW from *B. subtilis* (PDB: 5WUQ (12)). **C**: RpoE-RseA from *E. coli* (PDB: 1OR7 (14)). **D**: SigK-RskA from *M. tuberculosis* (PDB: 4NQW (13)).

Due to the diversity in ECF/ASDI quaternary structure, we here wondered whether there is a minimal contact interface conserved across all members of the ASDI family. To predict amino acid residues involved in such conserved contact interfaces, we turn to direct coupling analysis (DCA) - a bioinformatic method that exploits evolutionary co-variation to predict contacting residues (15). When two residues interact, mutations in one need to be compensated by changes in the second so as to preserve the interaction (15). The same mechanism also applies for indirect contacts; however, DCA is able to distinguish direct from indirect interactions and considers only the former for the calculation of their co-variation score (15). One of the highlights of DCA is that aside from stable conformations, it can also provide information on the transient, unstable conformations that occur during the dynamic process of interaction (16).

In this study we set out to provide the first phylogenetic classification of ASDI proteins, and reveal striking patterns of co-evolution between these regulators and their cognate ECF σ factors. For the ECF/ASDI interaction we used DCA to predict the residues that form the core ECF/ASDI contact. The arising sequence logos show divergent use of residues across ASDI groups, thus explaining the low binding affinity of non-cognate ECF/ASDI pairs from different groups. However, the predicted interaction partners in the fourth helix of ASDI and their respective counterparts are less conserved even within the ASDI groups. This might explain how ASDI proteins maintain binding specificity even within ASDI groups. These results allow a first, *in silico* assessment of potential cross-talk between two ECF/ASDI pairs without expensive *in vivo* testing, opening new ways to rationally design synthetic circuits using orthogonal ECF/ASDI pairs.

## Results

### ASDI retrieval and classification

We focused on the class I anti-σ factors (ASDI) as the main regulators of ECF σ factors, in order to gain a better understanding of their general binding mechanism to ECFs. Given that anti-σ factors are often co-encoded with their ECF targets (1, 2, 11, 17, 18), we first set out to collect ASDIs from the genetic neighborhood of the ECF coding sequences. To this end we focused on a set of 21,047 putative anti-σ factors identified during a recent classification effort of ECF σ factors by our group (3). To identify ASDI-containing proteins from this dataset, we used Hidden Markov Models (HMMs) developed from a small dataset of both zinc-binding and non-zinc binding ASDIs published earlier by Staroń and colleagues (2), see Methods for details. This step yielded 7,490 proteins, showing that ~36% of all putative anti-σ factors are ASDIs. In order to further expand the size of the ASDI sequence library, we built a new extended HMM from the ASDI domain of these sequences. We used this extended model to search for ASDIs in the genetic neighborhood of all classified ECFs identified in (3), using only the 33,843 ECFs from representative and reference organisms as labelled by the National Center for Biotechnology Information (NCBI). This yielded 11,939 proteins, from which we removed the ones with ASDI shorter than 50 amino acids, since these could be divergent class II anti-σ factors (19). The final number of ASDIs retrieved by this pipeline was 10,930, of which 10,806 have a non-redundant anti-σ domain. This shows that, on average, about one third (~32%) of the ECF σ factors contain a protein with an ASDI domain in their genetic neighborhood, suggesting that ASDIs are the most widespread regulators of ECF activity known to date. The average size of the ASDI domain was 101±33 (standard deviation) amino acids.

To gain an idea about the evolutionary history of these regulators, we classified them according to sequence similarity of their ASDI domain into 1,475 clusters of closely related sequences. For that, we used a divisive strategy, where the pool of sequences was subjected to a bisecting K-means clustering algorithm until the maximum k-tuple distance among sequences in the cluster was smaller than an empirical threshold of 0.6 (see Methods). Then, the consensus sequences of these clusters (referred to as subgroups hereafter) were hierarchically clustered into a phylogenetic tree (Fig. 2). A simple inspection of the ASDI tree shows that neighboring ASDI sequences on the tree also regulate σ factors from the same ECF group (Fig. 2, ring #2), supporting the notion that ECFs and ASDIs co-evolved. Given the similarity between ECF and ASDI classifications, we split the ASDI tree into monophyletic groups that regulate σ factors from the same ECF group (Fig. 2, ring #1). This split usually agrees with high bootstrap values (Fig. 2 and Fig. S1), suggesting that this definition of ASDI groups is robust to changes in the dataset. As a result, ASDI groups were named with “AS”, followed by a number dependent on the ECF group they are found in genomic proximity with. Even though ASDIs from the same clade of the ASDI tree are usually co-encoded with (and thus likely regulate) members of the same ECF group, some ASDI groups have slightly divergent features and are located in different clades of the ASDI tree. Two of these ASDI groups are for example AS19-1 and AS19-2, which regulate members of ECF19 (Fig. 2), but they are divergent in their ASDI helix 1 (consensus motif HTLAGAYALDAL in AS19-1 vs. HLDPDQLALLA in AS19-2) and helix 2 (consensus motif LDDERAAFERHL in AS19-1 vs. GEPLDADERAHL in AS19-2). Given that group AS19-2 is more closely related to AS27 than AS19-1, this suggests that these groups may have independently evolved the ability to bind to ECFs of group 19.

**Figure 2.**
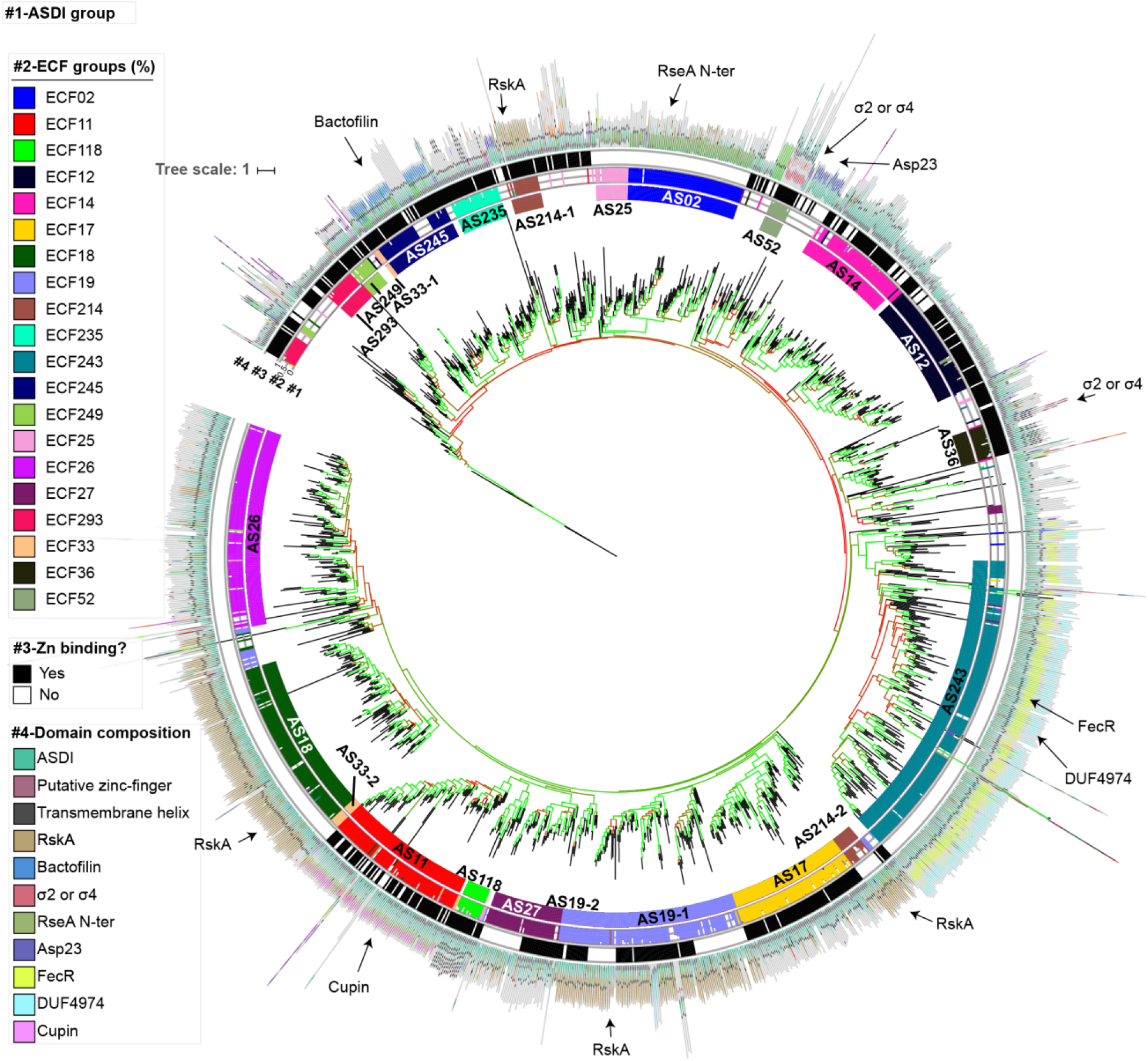
ASDI phylogenetic tree. Phylogenetic tree of the consensus sequences of subgroups of class I anti-σ factor domains. The tree is rooted at the sequence of the class II anti-σ factor CnrY, from *Cupriavidus metallidurans*., used as outlier. Branch length indicates evolutionary distance. Internal branch colors indicate bootstrap values, where 0% is red and 100% is green. Rings are explained as follows: #1) ASDI group defined in this work, #2) ECF group of the cognate ECFs encoded in the same genetic neighborhoods, #3) presence of Zn-binding motif, and #4) average domain composition of the anti-σ factors associated to each subgroup. The most important domains are explained in the legend.

ASDIs that regulate ECFs from the same subgroup are usually located together in the tree, but split into distinct ASDI subgroups (data not shown), probably due to the larger sequence diversity of anti-σ factors compared to ECFs. We observed that, even though there was a mixture of zinc-binding and non-zinc binding ASDIs in the input dataset (as indicated by the presence or absence of a ‘Hx_3_Cx_2_C’ motif), both types distribute across the ASDI tree, generating ASDI groups that are mixtures of zinc and non-zinc binding proteins, such as AS19-1 and AS27 (Fig. 2). Exceptions are groups AS33-1 and AS33-2, whose difference is the presence or absence of the zinc-binding domain, respectively (Fig. 2, ring #3).

Additionally, we predicted the mode number of transmembrane helices (TMHs) in the different ASDI subgroups using the consensus prediction from online TopCons (20). Most of the anti-σ factors (~65%) are predicted to contain at least one TMH, suggesting that they are bound to the membrane, while the remaining ones (35%) are likely soluble anti-σ factors. Although the whole dataset of ASDIs is composed of a similar amount of zinc- (~56%) and non-zinc-binding (~44%) proteins, we observed that amongst the soluble ASDIs there was an over-representation (~72%) of sequences with a zinc-binding motif. This is consistent with the notion that cytoplasmic ASDIs are often involved in sensing intracellular redox conditions (11, 21, 22). The membrane-bound anti-σ factors contained ~48% of sequences with zinc-binding motif, contrasting with earlier observations that membrane-bound anti-σ factors showed an under-representation of zinc-binding domains (11). However, the data in this earlier work was based on a must smaller sequence dataset of only 1266 sequences (11), suggesting that this apparent bias may have been due to random sampling of the sequenced genomes at the time. Our finding of an approximately equal distribution of zinc- and non-zinc-binding motifs indicates that sensing of internal redox conditions plays a lesser role in the membrane-bound ASDIs, and that the Zn-binding motif might take a more static, structural function as is the case for RsiW in *B. subtilis* (23).

If the Zn-binding motif does not play an active sensory role, the general notion is that the ASDI domains have associated with additional protein domains that allow stimulus perception and ultimately trigger anti-σ factor release (24, 25). To assess the conservation of additional protein domains, usually located C-terminal of the ASDI-domain, we scanned full-length class I anti-σ factors with Pfam 31.0 models (26) as well as the extended model of the ASDI domain. When indicating the positions of these domains in the different class I anti-σ factor subgroups (Fig. 2, ring #4), we found that the protein domains associated to ASDIs are typically well-conserved for ASDIs from the same group, but differ between groups. This suggests that ASDIs regulating members of the same ECF group are likely sensing similar input cues, either by binding directly to the triggering molecule or to other sensory proteins. The full list of ASDIs, together with their partner ECF and their ASDI group and subgroup can be found in Table S1.

Given the ample degree of correlation between ECF and ASDI classifications, we evaluated whether these families co-evolved. For this, we calculated the Pearson’s Correlation Coefficient (PCC) of the pairwise distance matrices of ASDIs and ECFs, as described by Goh and colleges (27), leading to a PCC of 0.82. In order to determine the significance of this correlation coefficient, we adopted the strategy by Dintner *et al.* (28) and included as negative controls RsbW-like anti-σ factors and RpoD-like σ factors, which do not interact with ECFs and ASDIs, respectively: RsbW is the anti-σ factor of the alternative σ factor σ^B^ and a protein kinase of the anti-anti-σ factor RsbV in *Bacillus subtilis* (29), while RpoD is the housekeeping σ factor of *Escherichia coli* (30). For these negative controls we obtained low PCCs (0.5 to 0.6), which are similar to the ones obtained by Dintner *et al.* for negative controls in bacterial two-component systems (28). This indicates that the PCC of 0.82 obtained for the correlation between ECFs and ASDIs is highly significant, showing that there has been strong co-evolution between these protein families (Table 1).

**Table 1.** **Pearson’s Correlation Coefficient** of the distances ECF and ASDI pairs in organisms that contain RsbW-like and RpoD-like proteins, used as negative controls for lack of correlation.

|  | ECFs | ASDIs | RsbW | RpoD |
| --- | --- | --- | --- | --- |
| ECFs | 1.00 |  |  |  |
| ASDIs | 0.82 | 1.00 |  |  |
| RsbW | 0.56 | 0.63 | 1.00 |  |
| RpoD | 0.48 | 0.50 | 0.67 | 1.00 |

### DCA predicts two main contact interfaces between ASDIs and ECFs

Given the variability in the binding conformations in the four published ECF/ASDI co-crystal structures, we next wondered whether there exist universally conserved “core binding interfaces” that are shared within the whole family of ASDI proteins, or whether the strong co-evolution between the protein families gave rise to fundamentally different binding conformations. To identify potentially conserved contact interfaces, we sought to exploit the co-evolutionary information between our ASDI dataset (above) and the ECF classification (3). Specifically, we aimed at predicting amino acid residues on ASDIs and ECFs that display significant co-variation, suggesting that they are in direct contact and that the mutation in one residue is balanced by a compensatory mutation in its binding partner. To this end we applied direct coupling analysis (DCA) (31) to the full set of ASDIs and their cognate ECFs (Table S1). The results of this analysis revealed a large amount of high DCA scores within the σ_2_ and σ_4_ domains of the ECF σ factor, and also connecting both σ domains (Fig. 3A). This pattern matches previous DCA results in ECF σ factors (32) and is indicative of the conserved secondary and tertiary structure on this family of proteins. We also observed high scores interconnecting helices 1, 2 and 3 of the ASDI, while helix 4 shows no strong DCA coupling scores with other parts of the ASDI domain (Fig. 3A). This agrees with the co-crystal structures of ECF/ASDI complexes (Fig. 1), where helices 1, 2 and 3 form a helix bundle, which is connected to helix 4 by a flexible linker (11–14). We then focused on the predictions that link ECFs and ASDIs since these are the ones responsible of ECF inhibition. At first glance, the contact map shows several high DCA scores linking the fourth helix of the ASDI with the σ_2_ domain (Fig. 3A). Under closer inspection, the top 14 inter-protein contact predictions (DCA score ≥ 0.255) are located in close proximity in most of the crystal structures (Fig. 3B and C). Of those, 12 are connecting the σ_2_ domain and helix 4 of the ASDI, and two (#10 and #11) connect a single residue of helix 1 of the ASDI to two residues located in the σ_4_ domain of the ECF (Fig. 3E). In the first case, the predicted contact area includes ECF regions 2.1 and 2.2, whose main function is binding to the clamp helices of the β’ subunit of the RNAP (33–35). Thus, it is likely that binding of ASDI’s helix 4 to this area prevents ECF binding to the RNAP core, hampering ECF-dependent transcription when the anti-σ factor in present. Instead, predictions #10 and #11 involve ECF helices 4.2 and 4.4, in two residues involved in the contact with the −35 element of the promoter (33, 36). Taken together, the presence of these strong co-evolutionary signals suggests that the majority of the 10,860 ASDI proteins establish contact to the ECF via these two binding interfaces, connecting the ASDI with both the σ_2_ and σ_4_ domains.

**Figure 3.**
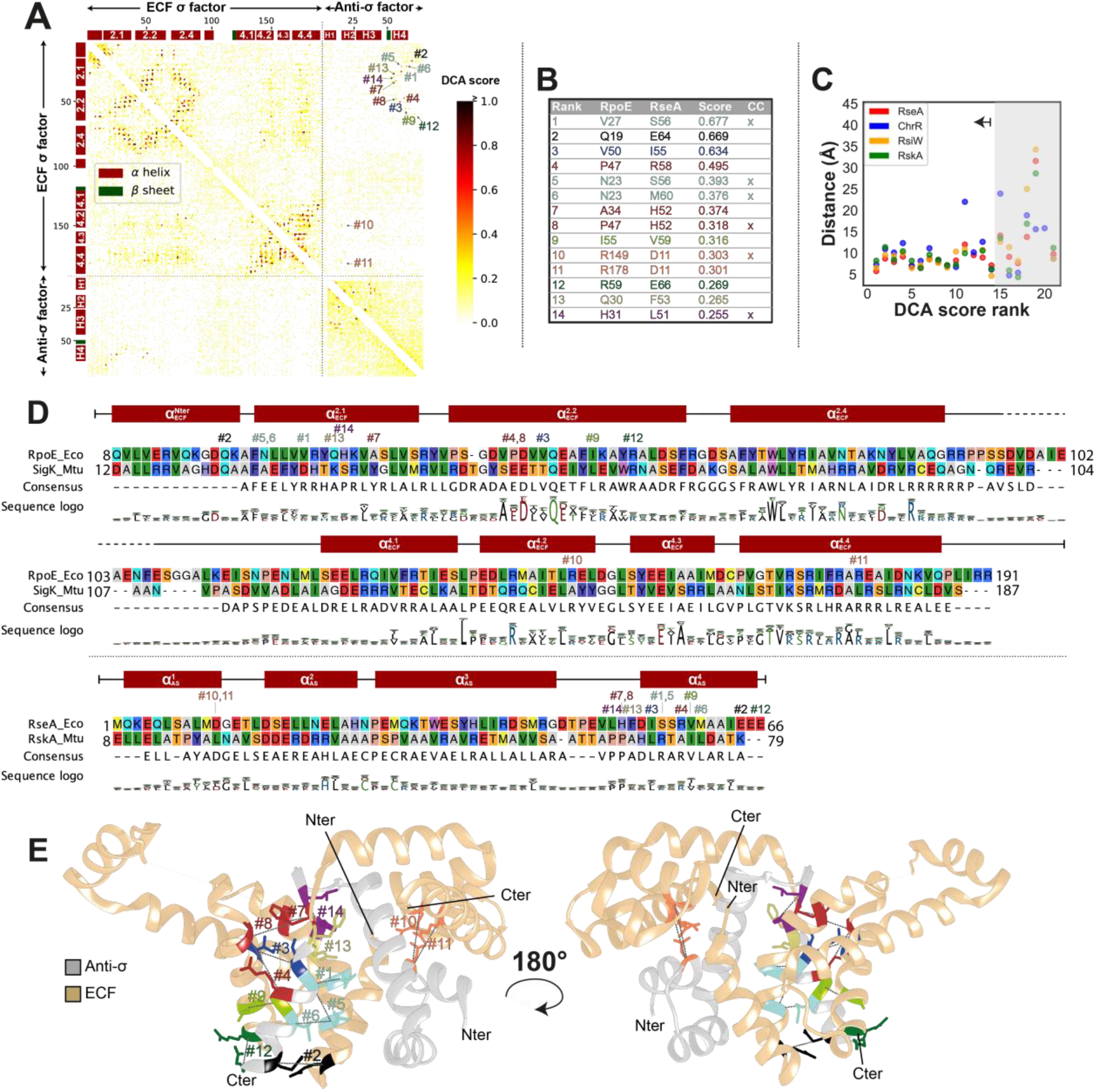
DCA results on the contact between ECFs and ASDIs. **A**: DCA contact map. Each axis represents the concatenated protein sequences of RpoE and RseA, from *E. coli,* used as reference for the amino acid labeling. High DCA scores, indicated by darker colors, correspond to residues with a high likelihood to bind *in vivo*. The largest 14 scores (DCA score ≥ 0.255) are marked in the heatmap and labelled according to their rank. **B**: Table of the 14 highest scoring DCA predictions, mapped to the amino acid coordinates of RpoE and RseA from *E. coli* are shown. The CC column indicates the DCA prediction that are also common contacts observed in the four crystal structures of ECFs/ASDIs, as derived by Voronoi tessellation (see Table 2). **C**: Scatter plot of the top 21 DCA predictions against the distance between the alpha carbons of the predicted contacts, as derived from the four structures of ECF/ASDI complexes (Fig. 1). The top 14 predictions are in close proximity in most of the 3D structures. Complexes are labeled after their anti-σ factor, where RseA corresponds to RpoE/RseA complex from *E. coli* (PDB: 1OR7 (14)), ChrR to SigE/ChrR from *R. sphaeroides* (PDB: 2Q1Z, (11)), RsiW to SigW/RsiW from *B. subtilis* (PDB: 5WUQ (12)) and RskA to SigK/RskA from *M. tuberculosis* (PDB: 4NQW (13)). **D**: Multiple-sequence alignment of two selected ECF/ASDI pairs, RpoE/RseA from *E. coli* and SigK/RskA, from *M. tuberculosis*. Labels of the top 14 contacts indicate their position. The presence of alpha helices and their names are depicted on top of the alignment. The sequence logo depicts the amino acid composition of the full ECF and ASDI alignments derived from 10,930 sequences, respectively. **E**: 3D depiction of the top 14 predictions in the structure of RpoE/RseA complex (PDB: 1OR7 (14)). ECF is colored in beige and anti-σ factor in gray. Predicted contacts are labeled according to their rank. N and C-termini from ECF and anti-σ factor are labelled.

However, although the top 14 DCA predictions connect residues located in close 3-dimensional proximity in most of the four resolved co-crystal structures of ECFs/ASDIs, only six are direct contacts in the four crystal structures (Fig. 3B). While the other 8 “close hits” could merely be false-positive predictions, it is tempting to speculate that these resides might form close contacts in other ECF/ASDI groups, which might take slightly different binding conformations from those captured by the four structures solved to date. Alternatively, these close hits may form transient contacts during the initial recognition between ASDI and ECF. Another observation was that 19 direct contacts that are shared between the four ECF/ASDI co-crystal structures were not predicted by DCA (Table 2), suggesting that DCA either fails to predict them, or that these contacts are less prevalent in the remainder of the ECF/ASDI protein families.

**Table 2.** Common contacts between ECFs and ASDIs found in the four co-crystal structures by Voronoi tessellation. The four crystal structures analyzed correspond to SigK/RskA from *M. tuberculosis* (PDB: 4NQW (13)), SigW/RsiW from *B. subtilis* (PDB: 5WUQ (12)), SigE/ChrR from *R. sphaeroides* (PDB: 2Q1Z (11)) and RpoE/RseA from *E. coli* (PDB: 1OR7 (14)). Coordinates of the different amino acids are shown in RpoE/RseA proteins. ECF and ASDI regions where the amino acids are located are shown. For simplicity, the σ_4_ domain is split into four subregions (σ_4.1_ - σ_4.4_) according to the presence of α helices. ‘H’ indicates α helix. The rank of the DCA prediction is displayed in the second last column when the interaction is predicted by DCA. If the residue is a SDP in the ASD, it is indicated in the last column.

| ECF residue<br>(mapped to<br>RpoE<sub>E. coli</sub>) | AS residue<br>(mapped to<br>RseA<sub>E. coli</sub>) | ECF<br>region | ASDI<br>region | DCA<br>prediction<br>(rank) | ASDI SDP ?<br>(Yes/No) |
| --- | --- | --- | --- | --- | --- |
| 22 | 59 | $\sigma_{2.1}$ | H4 | | N |
| 22 | 60 | $\sigma_{2.1}$ | H4 | | N |
| 22 | 63 | $\sigma_{2.1}$ | H4 | | N |
| 23 | 56 | $\sigma_{2.1}$ | H4 | 5 | Y |
| 23 | 60 | $\sigma_{2.1}$ | H4 | 6 | N |
| 26 | 55 | $\sigma_{2.1}$ | H4 | | N |
| 26 | 56 | $\sigma_{2.1}$ | H4 | | Y |
| 26 | 59 | $\sigma_{2.1}$ | H4 | | N |
| 27 | 56 | $\sigma_{2.1}$ | H4 | 1 | Y |
| 31 | 51 | $\sigma_{2.1}$ | Linker H3-H4 | 14 | N |
| 35 | 48 | $\sigma_{2.1}$ | Linker H3-H4 | | N |
| 47 | 52 | $\sigma_{2.2}$ | Linker H3-H4 | 8 | N |
| 47 | 55 | $\sigma_{2.2}$ | H4 | | N |
| 51 | 55 | $\sigma_{2.2}$ | H4 | | N |
| 51 | 58 | $\sigma_{2.2}$ | H4 | | N |
| 51 | 59 | $\sigma_{2.2}$ | H4 | | N |
| 54 | 59 | $\sigma_{2.2}$ | H4 | | N |
| 58 | 63 | $\sigma_{2.2}$ | H4 | | N |
| 131 | 42 | $\sigma_{4.1}$ | H3 | | N |
| 135 | 43 | $\sigma_{4.1}$ | H3 | | N |
| 149 | 11 | $\sigma_{4.2}$ | H1 | 10 | Y |
| 150 | 10 | $\sigma_{4.2}$ | H1 | | N |
| 150 | 40 | $\sigma_{4.2}$ | H3 | | N |
| 150 | 43 | $\sigma_{4.2}$ | H3 | | N |
| 151 | 43 | $\sigma_{4.2}$ | H3 | | N |

To obtain a better overview of the residues involved in the contact interfaces, we plotted the residues predicted by DCA - both in the ECF and in the ASDI - for the 12 largest ASDI groups with more than 100 sequences (Fig. 4). The resulting logos showed that contacts involving ASDI’s helix 1 and the σ_4_ domain (#10 and #11) are generally conserved within groups, but different between groups. Predictions #10 and #11 feature two main types of contacts, either a charged or a hydrophobic interaction (Fig. 4). This pattern is most evident for prediction #11, which tends to harbor a positive amino acid in the ECF (e.g. R178 in RpoE_*E.coli*_) and a negative residue in the ASDI (e.g. D11 in RseA_*E.coli*_), as found in groups ECF02, ECF12, ECF14, and ECF27. However, in some cases this is replaced by a hydrophobic contact, typically with leucine on both ECF and ASDI (e.g. L177 in SigK and L18 in RskA from *Mycobacterium tuberculosis*), as found in groups ECF17, ECF18 and ECF19. In contrast to these clear-cut contact motifs predicted for helix 1, residues in helix 4 of the ASDI (all predictions except #10, #11) exhibit a relatively limited conservation even within most of the ASDI groups (Fig. 4). Together with the observation that helix 4 of the ASDI holds most of the DCA predictions (Fig. 3D), this suggests that helix 4 is in charge of further determining the specificity of the ASDI, keeping them orthogonal from other ASDIs of the same group. Indeed, anti-σ factors that regulate ECFs from the same group have been found to be mostly orthogonal (37).

**Figure 4.**
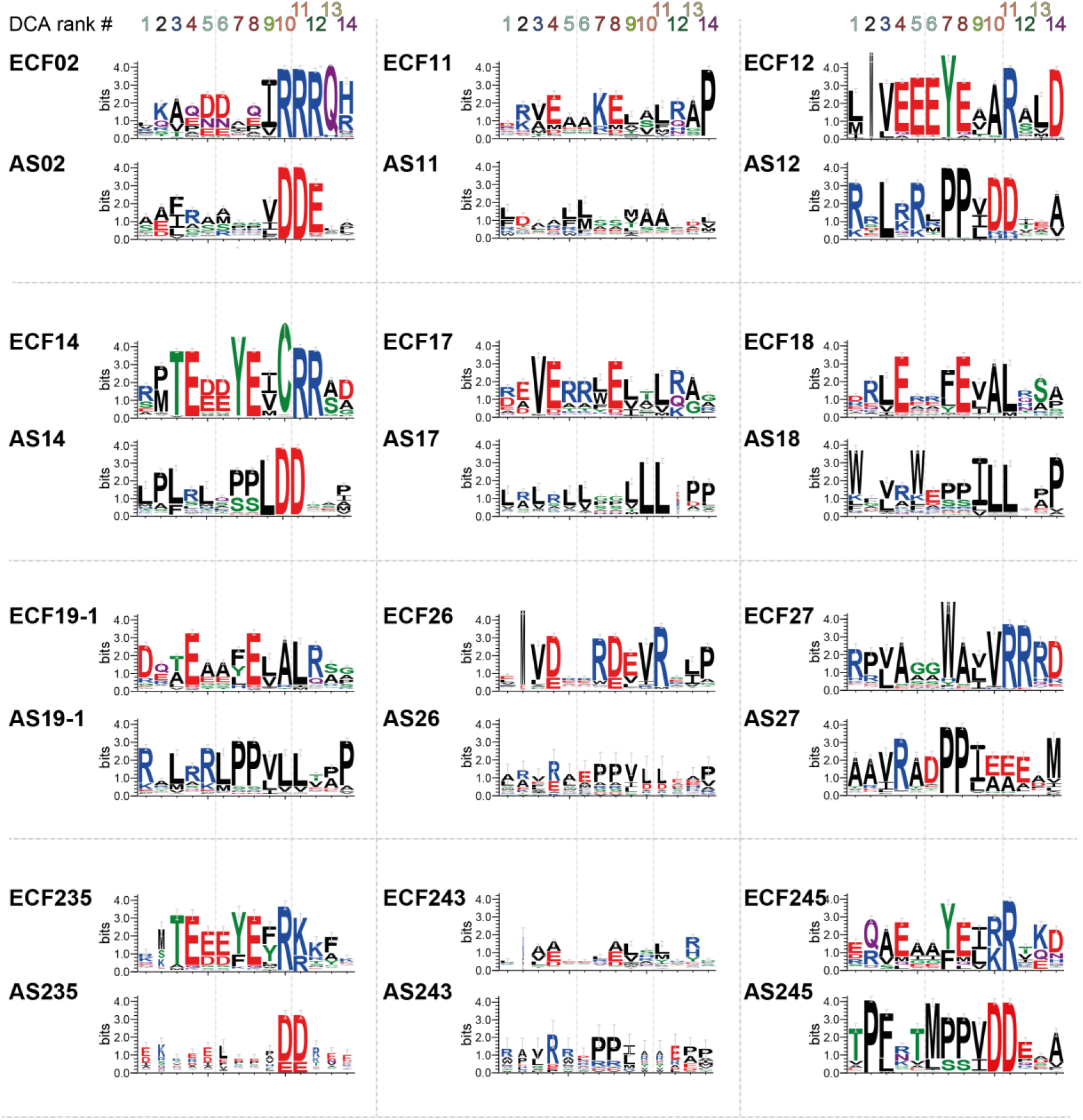
Sequence logos of the top 14 DCA predictions, computed for the 12 ASDI groups with more than 100 sequences. The sequence logos show the amino acid composition for the DCA-predicted contact points both for the ECF and anti-σ factor in each ECF/ASDI group. The contacts are ordered from left to right according to their DCA rank, as indicated on top.

### Specificity-determining positions of ASDI groups coincide with the predicted binding interfaces

Next, we asked whether the ASDI residues predicted to be in contact with the ECF are also key residues that determine the distinction between ASDI groups. If this was the case, it would suggest that ASDI groups would be primarily distinguished by their interaction with their respective ECF. Alternatively, if ASDI groups would primarily be determined by residues outside predicted contact interfaces, this would argue that interactions with potential ligands or intra-protein-interactions determine protein subfamilies (38). The presence of such group-specific amino residues - so-called Specificity Determining Positions (SDPs) - can be detected by S3det, a bioinformatic tool based in multiple correspondence analysis that finds residues associated to subfamilies of proteins (39). Using this tool, we predicted SDPs by comparing every pair of the 12 largest ASDI groups and taking only the highest scoring SDP prediction of every ASDI group into further consideration (see Methods). As a result, we identified five SDPs, named by running numbers (SDP#1 to SDP#5) from N- to C-terminus: two in helix 1, one in helix 3, one in helix 4 and the last one exclusively present in group AS243 (Fig. 5A). Proteins from group AS26 did not hold any prediction, since they do not fit well into the multiple sequence alignment of the full ASDI dataset - probably due to extensive differences at the sequence level (cf. Fig. 4). Similarly, AS243’s SDP corresponds almost exclusively to a gapped position in the alignment with the rest of the groups (Fig. 5C #5). These differences at sequence level might reflect functional differences between standard ASDIs and ASDIs from groups 243 and 26. In favor of this hypothesis, one member of AS243, FecR from *E. coli*, is distinguished from other non-AS243 ASDIs in that its 59 N-terminal amino acids are required for ECF activity (40). Interestingly, all predicted SDPs are part of the contact interfaces with the ECF in the existing crystal structures (Fig. 5B, Fig. S2). Conserved position D11 in RseA_*E.coli*_, predicted by DCA (Fig. 3B, #10 and #11), was part of the predicted SDPs (Fig. 5A, SDP#2). Yet another SDP, V27 in helix 4 (Fig. 5A, SDP#4), was predicted by DCA (Fig. 3B, #1 and #5). Predictions SDP#1 and SDP#3 connect S7 in helix 1 and Y36 in helix 3 in RseA_*E.coli*_ to the σ_4_ domain, usually in its last helix (Fig. 5B, Fig. S2). Interestingly, SDPs #1, 2 and 3 form a cluster of interactions with the same area of the ECF, which usually corresponds to the last helix of the σ_4_ domain, except in SigE/ChrR structure, where the contact appears before this area (Fig. 5B, Fig. S2). Thus, besides some exceptions in groups AS26 and AS243, these results suggest that the main characteristic that discriminates between ASDI groups is their ability to interact with the σ factors within their cognate ECF groups.

**Figure 5.**
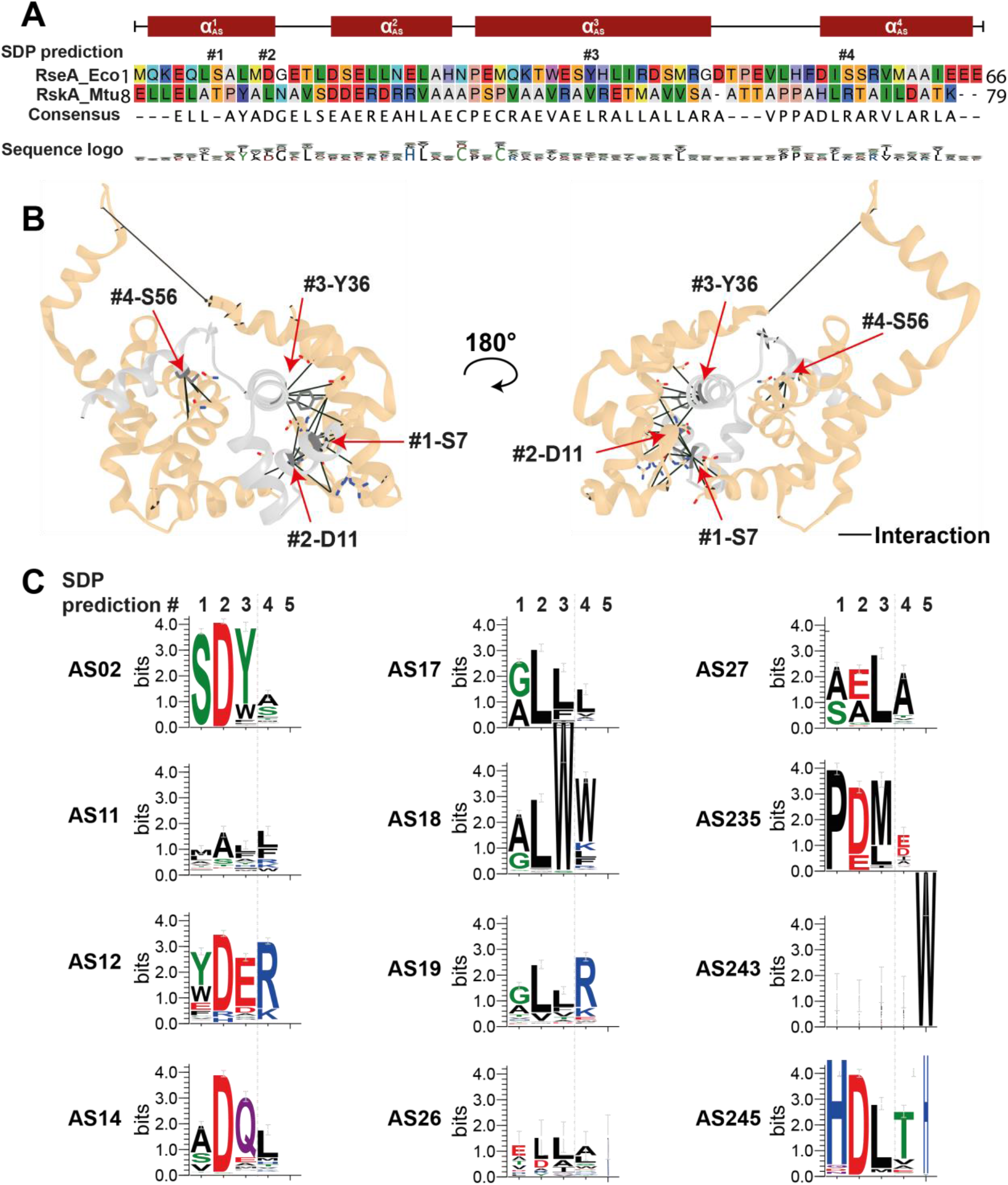
Description of the specificity-determining positions (SDPs) that distinguish different ASDI groups. **A**: Multiple-sequence alignment of the anti-σ factors RseA from *E. coli* and RskA from *M. tuberculosis* showing the position of the SDPs, labelled with numbers according to sequence position. Alpha helices and their names are indicated with red boxes on the ASDI sequences. The sequence logo shows the amino acid composition of the full ASDI alignment. **B**: Structure of the RpoE/RseA complex from *E. coli* (PDB: 1OR7 (14)). SDPs are labeled as in **A** and their contacts with the ECF are shown by lines. **C**: Logo of SDPs in every ASDI group with more than 100 proteins. Positions are labelled as in **A**.

Given that these residues are conserved within phylogenetic ASDI groups, face the ECF in the solved ECF/ASDI crystal structures and feature different amino acids in different groups, it is likely that they take part in determining specificity towards the target ECF. This is supported that the fact that most of these SDPs are also DCA predictions (Table 2).

## Discussion

In this study, we used a computational approach to study how class I anti-σ factor family interact with their cognate ECF σ factors. Based on the similarity between ECF and ASDI phylogenies, we showed that these protein families have co-evolved - likely because they are in direct contact with each other - and exploited this co-evolution to predict two conserved binding interfaces for the ASDI/ECF interaction. Although previous work provided insight in the co-crystal structures of individual ASDI/ECF pairs, the present work puts these case-studies into a broader, evolutionary perspective, by providing the first phylogenetic classification of the class I anti-σ factor protein family. Interestingly, within the resulting AS groups - solely defined by the sequence of their ASDI domain - we observed a striking conservation of the fused protein domains. Compared to early work by Campbell and colleagues (11), the explosion in sequenced genomes in recent years allowed us to expand the ASDI dataset from 1266 to more than 10,000 putative ECF/ASDI pairs from NCBI reference genomes, providing a more comprehensive and phylogenetically balanced overview on the diversity of these proteins. In agreement with Campbell *et al.* we found that about one third (~32%) of all ECFs are genomically associated with, and thus likely regulated by ASDIs. Yet, our expanded ASDI library showed important differences compared to previous work in that, (i) we find more ASDIs containing a zinc-binding motif (~56% compared to ~38% (11)), (ii) we find more cytoplasmic anti-σ factors (~35% compared to ~28% (11)), (iii) cytoplasmic anti-σ factors are still overrepresented in zinc-binding motifs, but to a smaller extent (~72% of the soluble anti-σ factors are zinc-binding in our dataset compared to 92% in (11)), and (iv) membrane-bound ASDIs are not underrepresented in zinc-binding motifs as suggested in (11), with about half of the proteins (~48%) being zinc binding anti-σ factors. These data suggest that ASDIs are more diverse than previously thought, and argues against a functional role of the zinc-binding domain exclusively in soluble anti-σ factors. This is supported by the ASDI phylogenetic tree (Fig. 2), where zinc and non-zinc binding ASDI groups are mixed across the tree and sometimes even within the same group, as in the case of AS27, and AS19-1. In these mixed zinc and non-zinc binding groups, this suggests that the zinc-binding motif may play a structural instead of a sensory role, as shown for RsiW from *B. subtilis* (group AS245) (12).

Our analysis of DCA predictions and SDPs show that there exists a conserved, dual binding interface, with ASDI’s helix 1 binding to the σ_4_ domain and ASDI’s helix 4 binding to the σ_2_ domain. These results agree with crystal structures of ECF/ASDI complexes (11–14) and suggest that the contacts seen in these few examples are indeed realized across the full ECF/ASDI families. Further, our results suggest that ASDI’s helix 2 is not critical for ECF binding but is important for ASDI tertiary structure. ASDI′s helix 3, which is located between ECF’s σ_2_ and σ_4_ domains in three out of four structures (11, 13, 14), harbors a SDP involved in the interaction with σ_4_ domain, in similar residues as contacted by the prediction on helix 1. This modularity of the ASDI interaction is reflected in the function of the ECF residues involved in the predictions. Contacted residues in regions 2.1 and 2.2 are mostly involved in the contact with the clamp helices of the β’ subunit of the RNAP (33, 35), whereas predicted contacts in σ_4_ are part of the contact interface with the −35 element of the promoter (33, 36).

The analysis of the DCA predictions revealed a different degree of conservation across ASDI groups, with the residues that take part in contacts between ASDI’s helix 1 and ECF’s σ_4_ (DCA predictions #10 and #11) being conserved for most of the ECF and ASDI phylogenetic groups. Interestingly, this area, which connects D11 on the ASDI to R149 and R178 on the ECF (RseA/RpoE_*E.coli*_ coordinates) bears two main types of interactions, that is, hydrophobic, which usually features leucine in both ECF and ASDI (Fig. 4, groups AS17, AS18 and AS19-1), or charged, usually featuring arginine in the ECF side and aspartate in the ASDI side (Fig. 4, groups AS02, AS12 and AS14, among others). Random mutagenesis in RseA_*E.coli*_ (group AS02) showed that a single amino acid mutation of D11 to histidine completely inhibits RseA_*E.coli*_ activity (41), confirming the key role of this contact. Given their group-specific conservation and the striking polarity differences between the two binding types, we speculate that D11 defines coarse-grained specificity of ASDIs for ECFs of the same binding type, usually found in the same phylogenetic group. However, ASDIs are usually specific to their own target ECF and do not usually crosstalk with members of the same group (37), indicating that there are more sources of specificity in residues that are not conserved in groups. One potential source of this specificity are the residues predicted by DCA in helix 4. These residues are generally not conserved within groups (Fig. 4) and bind the σ_2_ domain in all the solved crystal structures of ASDI/ECF complexes (11–14). This lack of major conservation is extended to the predicted contacts on the ECF side, which are generally in charge of binding to the β’ subunit of the RNAP.

### Generality of the dual binding interface in other σ/anti-σ interactions?

Paget classified anti-σ factors into two types, the ones that insert between σ_2_ and σ_4_ (RseA, RskA and ChrR) and the ones that wrap around these domains (RsiW) (42). Our data shows that despite these differences in binding topology, both types of ASDIs contact the two main binding interfaces described here. Moreover, a similar binding mode can be observed in the crystal structures of the ECF CnrH in complex with the class II anti-σ factor CnrY, from *Cupriavidus metallidurans* (43). The two α helices of CnrY wrap around CnrH in a conformation where CnrY’s first α helix mimics the function of ASDI’s first helix and binds to σ_4_ domain, and CnrY’s second and last α helix binds to σ_2_ domain in a similar manner as ASDI’s fourth helix. The only crystal structure of a member of the ASDIII class of anti-σ factors, BldN, in complex with the ECF σ factor RsbN from *Streptomyces venezuelae* (44) also shows this dual binding mode. In this case, the first and second α helices of BldN bind to the σ_4_ domain, whereas its third and last α helix binds to the regions 2.1 and 2.2 of a different RsbN molecule, similarly to ASDI’s forth helix (44). The similarity of the binding between the three types of ECF anti-σ factors is striking and contrasts with their low level of sequence similarity, which is limited to ~11% for RseA/BldN and ~3% for RseA/CnrY (using global pairwise alignments calculated by Needleman-Wunsch algorithm implemented at EBI (45)). This explains why, even though the same regions of the anti-σ factor interact with a similar area of the ECF in the three types of ECF anti-σ factors, the specific residues that carry out the interaction with the ECF may differ between ASD types. It is unclear why bacteria need at least three types of ASDs. On one hand, different ASDs may provide extra specificity to ECF inhibition, which could help to reduce the apparent tendency to cross-talk of anti-σ factors (46). On the other hand, the three types of ASDs could have emerged from different proteins and optimized their ECF inhibition by blocking the same ECF regions through convergent evolution. Future analysis that include all the ASDs known to date could help in understanding their evolution.

Interestingly, dual binding interfaces between σ and anti-σ factor extend beyond ECF σ factors. For instance, in *E. coli* the anti-σ factor FliM of the class 3 σ factor FliA (containing a σ_3_ domain) also targets σ_2_ and σ_4_ regions with two different areas of the protein (47). However, the FliM inhibitory contacts are inverted relative to ECF anti-σ factors: FliM is composed of four α helices, of which the first and second bind to the surface of the σ_2_ domain, similarly to the fourth helix of ASDIs. In FliM, the third and fourth helices are the ones that bind to σ_4_ (47), similarly to the first helix of ASDIs. Interestingly, FliM does not bind to FliA’s σ_3_ domain, strengthening the idea that the blockage of both σ_2_ and σ_4_ are the core of σ factor inhibition. Whether this is also the case for housekeeping σs and their anti-σs remains to be seen, as to date only the interaction between the anti-σ factor Rsd and a truncated version of RpoD (only containing the σ_4_ domain) was studied in *E. coli* (48, 49). Thus, even though the present analysis was restricted to the interaction between ASDIs and ECFs, we suggest that the dual inhibition of RNAP- and DNA-binding interfaces is likely a universal feature of other anti-σ factors, preventing formation of non-functional trimeric complexes between σ/anti-σ factors and RNAP or DNA.

## Acknowledgements

This work was supported by a grant from the ERA-SynBio program via the Federal Ministry of Education and Research (Germany; grant 031L0010B) and the LOEWE program of the State of Hesse (Germany).

## Methods

### General bioinformatic tools

Generally, Multiple Sequence Alignments (MSAs) were generated by Clustal Omega 1.2.3. with options --iter=2 and --max-guidetree-iterations=1 and manually curated (50). However, UPP (51) (default options) was used for alignments subjected to DCA or to S3det, since they require stable columns of equivalent residues with few gaps. Hidden Markov Models (HMMs) were built using *hmmbuild* function and used for scanning libraries using *hmmscan* function, both from HMMER suite 3.1b2 (52) and both with default parameters. For the extraction of the amino acid residue interactions between ECF and ASDI from co-crystal structures, we used Voronoi tessellation as implemented in Voronota version 1.19 (53). Protein structures were visualized using UCSF Chimera version 1.10.2 (54).

### ASDI extraction

ASDIs were extracted from the genetic neighborhood (±10 coding sequences) of a library of 46,293 ECF σ factors in their most recent classification (3). In order to minimize taxonomic bias, these ECFs were extracted from organisms tagged as representative or reference species by NCBI (https://www.ncbi.nlm.nih.gov/refseq/about/prokaryotes/), using only RefSeq entries when both RefSeq and GenBank records are available for the same genome. To identify ASDI domain-containing proteins, we first used two HMMs, one built from the zinc-binding and another from the non-zinc-binding anti-σ factors from Staroń *et al.* (2). We selected the optimal bit score threshold for the retrieval of new ASDIs for each HMM by optimizing a Receiver Operator Characteristic (ROC) curve using the function *roc_courve* from *sklearn.metrics* (55). Proteins that were used for the construction of each model were used as positive controls and the remaining, non-ASDI anti-σ factors from Staroń *et al.* (2) as negative controls. The resulting bit score thresholds, 0.4 for non-zinc binding and 14.2 for zinc-binding models, were applied for the extraction of ASDIs from the set of putative anti-σ factors from (3). This resulted in 7,490 ASDIs, which were subsequently used for the construction of an extended HMM of the ASDI family. The thresholding bit-score that best separates real ASDIs from other proteins was optimized using a ROC curve as described above, resulting in a bit-score threshold of 0.2. We used the extended HMM to look for further members of the ASDI family in the genetic neighborhood of ECFs (±10 coding sequences) from (3). In order to lessen the bias towards frequently sequenced organisms, we only included proteins from representative or reference genomes as labelled by NCBI (https://www.ncbi.nlm.nih.gov/refseq/about/prokaryotes/), using only RefSeq entries when both RefSeq and GenBank records are available for the same genome. This yielded 11,939 putative ASDI-containing proteins. We further curated these data removing proteins with anti-σ domains shorter than 50 amino acids, since these could be anti-σ factors of class II (19). The area of the anti-σ domain was defined as the envelope region of the highest scoring hit of the extended HMM, discarding areas that are part of the transmembrane helices or extracellular. This resulted in 10,930 ASDIs, with an average length of 101± 33 (standard deviation) amino acids.

### Clustering of ASDIs

We clustered ASDIs according to amino acid sequence similarity. Given the large number of proteins, we first grouped them into clusters or closely related sequences, the so-called subgroups. These were built with a divisive strategy, where proteins were subjected to a bisecting K-means clustering approach until the maximum k-tuple distance between any protein of the cluster is smaller than 0.6, as measured by Clustal Omega with --distmat-out --full and --full-iter flags (50, 56). Bisecting K-means was implemented using *KMeans* function from *sklearn.cluster* module (55). The 3,790 proteins that did not enter into any subgroup were left ungrouped. Thanks to this grouping it was easier to see subgroups that may contain outliers that passed the HMM threshold, but do not likely display anti-σ factor activity. In order to distinguish and discard these outliers from our clustering, we assessed the presence of Pfam domains (Pfam 31.0 (26)) in the anti-σ factors from each subgroup. We discarded 132 subgroups (606 proteins) where the Pfam domains indicated an unlikely anti-σ factor function (data not shown). In summary, the resulting 1,475 subgroups defined during this process contained 6,534 proteins (~60% of the starting ASDIs), with a median group size of 3 proteins and a standard deviation of 6.17 proteins. Given the low size of proteins in each subgroup, we further clustered the manually curated alignment of the consensus sequences of each subgroup, into a maximum-likelihood phylogenetic tree using IQ-TREE version 1.5.5 (57) with 1000 ultrafast bootstraps. As an outgroup of this tree, we included the anti-σ factor class II CnrY, from *Cupriavidus metallidurans.* The resulting tree was visualized in iTOL (58) and split into monophyletic ASDI groups according to the ECF group of their cognate partner. With this strategy we defined 23 ASDI groups, of which 12 contain more than 100 proteins.

The presence of a zinc-binding domain was assumed in ASDIs with a Hx_3_Cx_2_C sequence signature that expands over helix 2 and helix 3. Presence of transmembrane helices was assessed using the consensus prediction from online TopCons (20). The mode number of transmembrane helices was considered in order to plot the transmembrane helices for a whole subgroup of class I anti-σ factors. In this way we avoid biases caused by the extreme large number of transmembrane helices in long, divergent proteins. The position of these helices for plotting was calculated according to the average start and end positions over the anti-σ factors in a subgroup. Similarly, the position of the ASDI domain across anti-σ factors from the same subgroup was calculated according to the average start and end positions of the envelope region of the lowest E-value match to the extended HMM of the ASDI family, using hmmscan for HMMER suite 3.1b2 (52). The presence of other Pfam domains in full-length class I anti-σ factors was evaluated using hmmscan function from HMMER suite 3.1b2 (52) with the library of HMMs from Pfam 31.0 (26). Pfam domains present in certain position of the MSA of the full-length anti-σ factors in more than 50% of the members of a subgroup were plotted in the ASDI tree.

### ASDI/ECF co-evolution

In order to evaluate the co-evolution of ECFs and ASDIs, we calculated the Pearson correlation coefficient (PCC) of the distances between cognate pairs of proteins, as introduced by Goh *et al.* (27). The significance of this PCC was evaluated similar to (28). For this purpose, the PCCs between ASDIs, ECFs and of two extra families of proteins that did not co-evolve and/or interact with ECFs or ASDIs were evaluated as negative controls. In our case, these negative controls were homologs of *E. coli*’s housekeeping σ factor σ^70^ (RefSeq: NP_417539.1) and of *Bacillus subtilis*’ anti-σ factor RsbW (RefSeq: WP_061902497), since proteins for these types have never been described to interact with ASDIs nor ECFs, respectively. We extracted proteins from these types using online HMMER (52) with parameters -E 1 --domE 1 --incE 0.01 --incdomE 0.03 --mx BLOSUM62 --pextend 0.4 --popen 0.02 -- seqdb uniprotrefprot, and mapped the hit IDs from UniProt to GenBank using the UniProt’s ID conversion tool (59). A total of 409 genomes contained the four protein families, this is ECFs, ASDIs, RsbW and RpoD. For each organism, we selected one of the ECF-AS factor pairs and one homolog of RsbW and RpoD. These proteins had a taxonomic diverse origin, with 39% of the proteins from Firmicutes, 28% from Actinobacteria, 11% from Cyanobacteria and the rest from other eight bacterial phyla. We calculated the pairwise distance for each protein family using Clustal Omega with -full and --distmat-out flags (50). The PCC was calculated from the flattened distance matrices using *pearsonr* function from Python’s *scipy.stats* resource (60).

### Direct Coupling Analysis (DCA)

DCA was applied to the 10,930 putative ASDIs extracted during this work (Table S1). ASDI and their cognate ECF partners were aligned independently using UPP (51) with default parameters, and the resulting alignments were concatenated. Gaussian DCA with default parameters (61) was performed on this alignment (N=275, M=10,934, Meff=965.52, theta=0.46). The top DCA predictions were mapped into the crystal structures of RpoE/RseA from *Escherichia coli* (AS02, PDB: 1OR7 (14)), SigE/ChrR from *Rhodobacter sphaeroides* (AS11, PDB: 2Q1Z (11)), SigK/RskA from *Mycobacterium tuberculosis* (AS19-1, PDB: 4NQW (13)) and SigW/RsiW from *Bacillus subtilis* (AS245, PDB: 5WUQ (12)). Distances between predictions were calculated using Bio.PDB module (62, 63) and Chimera (54). The 14 predictions that connected residues in close proximity (<15Å) in most of the structures were considered true interactions.

### Specificity Determining Positions (SDPs)

SDPs were calculated with S3det (39) on the 12 ASDI groups with more than 100 proteins, and on their cognate ECFs. Aligned ASDI (or ECF) proteins were extracted from the MSA used for DCA so as to preserve the same positional mapping. S3det was executed on every pair of ASDI (or ECF) groups, resulting in a set of ranked SDP predictions for every pair of groups. We scored the SDPs associated to every group as the sum of the inverse of their ranks across the different S3det runs with contribution of the group. The highest scoring SDP for every group was considered positive, resulting in five SDPs.

---

**Figure S1.**
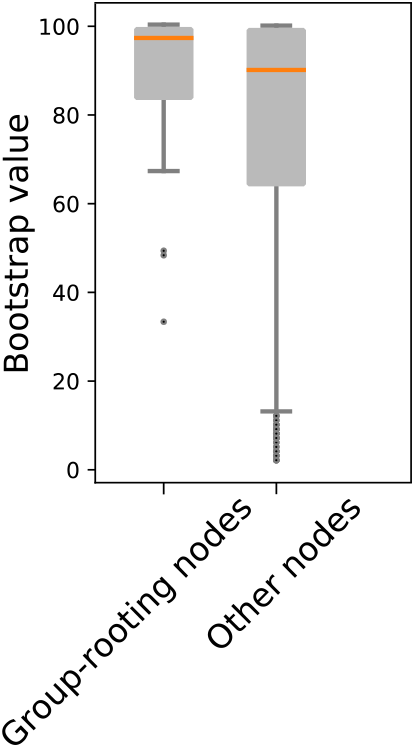
Bootstrap value distribution of the different branches of the ASDI tree (Fig. 2). ASDI rooting nodes, which are in the base of ASDI trees, have generally higher bootstrap values.

**Figure S2.**
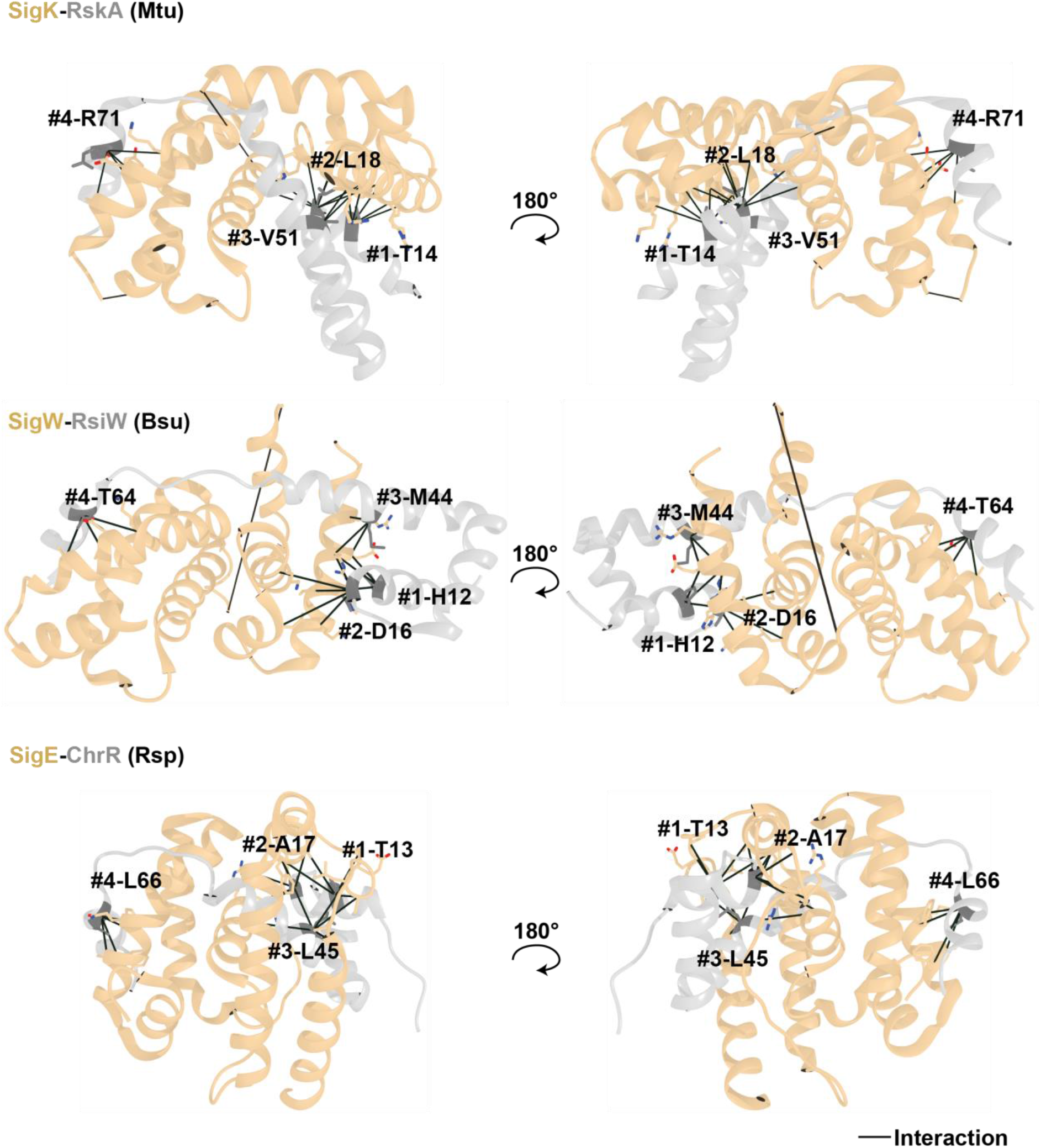
ASDI specificity-determining positions plotted in the structure of ECF/ASDI complexes. ECFs are colored in beige and anti-σ factors in gray; SDPs are colored in green and labelled with their identifier as in Fig. 5. SigK/RskA complex is present in *M. tuberculosis* (Mtu, PDB: 4NQW (13)), SigW/RsiW in *B. subtilis* (Bsu, PDB: 5WUQ (12)) and SigE/ChrR in *R. sphaeroides* (Rsp, PDB: 2Q1Z (11)). Contacts with the ECF are represented by connector lines. A similar representation is available for RpoE/RseA pair in Fig. 5B.

**Table S1. Full list of class I anti-σ factors used during this study.** Anti-σ factors are associated to their ASDI group and subgroup, their protein sequence, the ECF from whose genetic neighborhood it was extracted, the ECF’s group and subgroup, their organism of origin and NCBI identifiers of both proteins.

